# Generation of inducible SMARCAL1 knock-down iPSC to model severe Schimke immune-osseous dysplasia reveals a link between replication stress and altered expression of master differentiation genes

**DOI:** 10.1101/546093

**Authors:** Giusj Monia Pugliese, Federico Salaris, Valentina Palermo, Veronica Marabitti, Nicolò Morina, Alessandro Rosa, Annapaola Franchitto, Pietro Pichierri

## Abstract

The Schimke immuno-osseous dysplasia is an autosomal recessive genetic osteochondrodysplasia characterized by dysmorphism, spondyloepiphyseal dysplasia, nephrotic syndrome and frequently T cell immunodeficiency. Several hypotheses have been proposed to explain pathophysiology of the disease, however, the mechanism by which SMARCAL1 mutations cause the syndrome is elusive. Indeed, animal models of the disease are absent or useless to provide insight into the disease mechanism, since they do not recapitulate the phenotype. We generated a conditional knockdown model of SMARCAL1 in iPSCs to mimic conditions of cells with severe form the disease. Here, we characterize this model for the presence of phenotype linked to the replication caretaker role of SMARCAL1 using multiple cellular endpoints. Our data show that conditional knockdown of SMARCAL1 in human iPSCs induces replication-dependent and chronic accumulation of DNA damage triggering the DNA damage response. Furthermore, they indicate that accumulation of DNA damage and activation of the DNA damage response correlates with increased levels of R-loops and replication-transcription interference. Finally, we provide data showing that, in SMARCAL1-deficient iPSCs, DNA damage response can be maintained active also after differentiation, possibly contributing to the observed altered expression of a subset of germ layer-specific master genes. In conclusion, our conditional SMARCAL1 iPSCs may represent a powerful model where studying pathogenetic mechanisms of severe Schimke immuno-osseous dysplasia, thus overcoming the reported inability of different model systems to recapitulate the disease.

## INTRODUCTION

The Schimke immuno-osseous dysplasia (SIOD) is an autosomal recessive genetic osteochondrodysplasia characterized by dysmorphism, spondyloepiphyseal dysplasia, nephrotic syndrome and frequently T cell immunodeficiency ^1–3^. Patients usually suffer of other less penetrant features and, depending on the severity of the disease, they can undergo to premature death in the childhood or early adolescence ^3^. The disease is caused by bi-allelic mutations in the SMARCAL1 gene ^4^. *SMARCAL1* encodes for a protein homologous to the SNF2 family of chromatin remodeling factors, however, recent works firmly demonstrated that SMARCAL1 is not involved in chromatin remodeling and transcriptional regulation, but rather in the processing of DNA structures at replication forks to promote formation of replication intermediates through its ATP-driven strand-annealing activity ^5,6^. Many mutations in the SMARCAL1 gene have been identified, ranging from frameshift and deletions that generally lead to protein loss, to missense mutations that differently affect expression, activity, stability and localization of the protein ^1,7^.

Basing on the pathophysiology of the disease several hypotheses have been proposed ^1,8^, however, the mechanism by which SMARCAL1 mutations cause SIOD are completely unknown. The recent demonstration that SMARCAL1 is critical to the response to perturbed replication and that its loss or impaired activity hampers recovery from replication stress and determines DNA damage formation, challenged the canon for SIOD hypothetical molecular pathology from transcriptional regulation to DNA damage prevention. Thus, it is tempting to speculate that, similarly to Seckel syndrome, Werner’s syndrome and other genetic conditions caused by loss of genome caretaker proteins, SIOD may be generated by an accumulation of DNA damage and impaired proliferation or development that could follow.

Interestingly, SIOD patients bearing distinct SMARCAL1 mutations show a different degree of disease severity ^7^. Thus, a phenotype-genotype correlation might exist although difficult to ascertain. Indeed, mutations resulting in the almost complete loss of protein are associated to severe SIOD. By contrast, mutations that similarly affect SMARCAL1 ATPase activity give raise to both severe or mild SIOD, arguing for the existence of genetic factors that can modulate disease phenotypes or of additional ATPase-independent SMARCAL1 functions that are affected by missense mutations ^7–9^.

Unfortunately, models of SIOD are absent or were useless to provide insight into the disease mechanism. Indeed, deletion of SMARCAL1 in mice or fruit flies fails to recapitulate the disease phenotype ^9^. Only a study from zebrafish evidenced cell proliferation and developmental defects upon deletion of the smarcal1 orthologue ^10^, suggesting that loss of SMARCAL1 could affect proliferation and development in humans too. Thus, although likely to exist, the correlation between SMARCAL1 mutations, replication stress, DNA damage formation, defects in proliferation and impaired development in SIOD pathogenesis is yet completely unexplored, largely because of the absence of useful models of the disease.

Induced Pluripotent Stem Cells (iPSCs) are very informative on the very first stages of development. Such model system is very useful for the identification of early events associated to disease pathophysiology. Moreover, it is genetically amenable and provides cell types for drug screening.

Here, we generated a model to study severe SIOD generating iPSCs in which expression of SMARCAL1 could be downregulated through a Tet-ON-regulated RNAi system. Using this cell model, we demonstrated that depletion of SMARCAL1 resulted in reduced proliferation, accumulation of DNA damage, replication defects and DNA damage response overactivation. Moreover, our data show that the most striking phenotypes are correlated with increased R-loop accumulation and can be reversed preventing replication-transcription interference. Most importantly, using our iPSC cell model of severe SIOD, we show that replication-related DNA damage persists also in differentiated cells and that loss of SMARCAL1 affects expression of a subset of germ layer-specific marker genes.

## RESULTS

### Generation and characterization of inducible SMARCAL1 knockdown iPSCs

To obtain an inducible model of severe SIOD, we expressed an shSMARCAL1 cassette under the control of a Tet-ON promoter through lentiviral transduction in the well-characterized normal iPS cell line WT I ^11^ (Fig. 1A). Low-passage iPSC were infected with the Tet-ON-shSMARCAL1 virus at 0.5 of MOI by spinfection, selected and tested for the knockdown efficiency by Western blotting. As shown in Figure 1B, culture of inducible SMARCAL1 knockdown (iSML1) iPSC with doxycycline (DOX) for 48h resulted in less than 13% of total SMARCAL1. Western blotting analysis of the SMARCAL1 level after 7 or 14 days of continuing growth in DOX revealed that the high knockdown efficiency was stable over time in the iSML iPSC (Fig. 1C).

**Figure 1.**
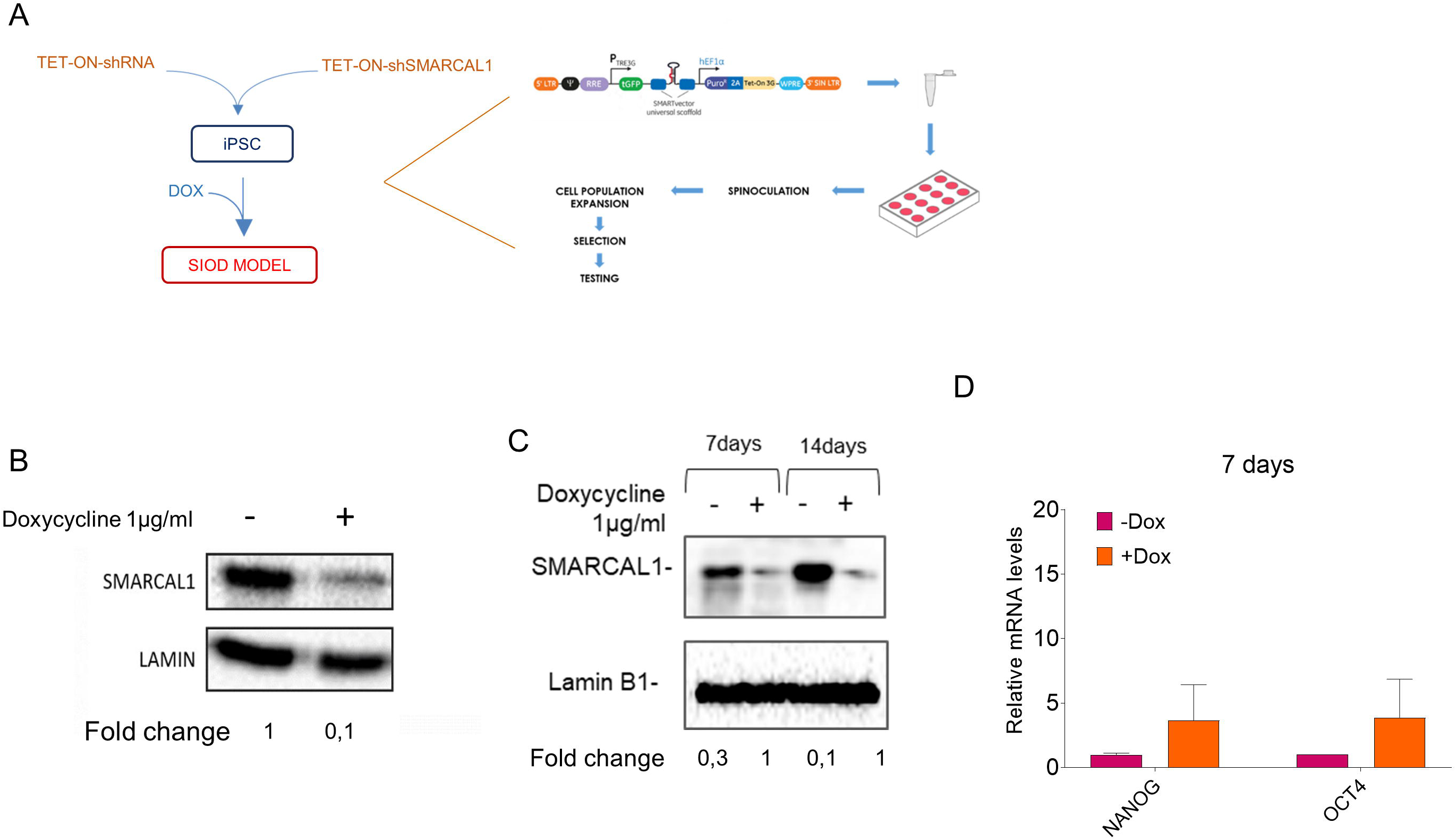
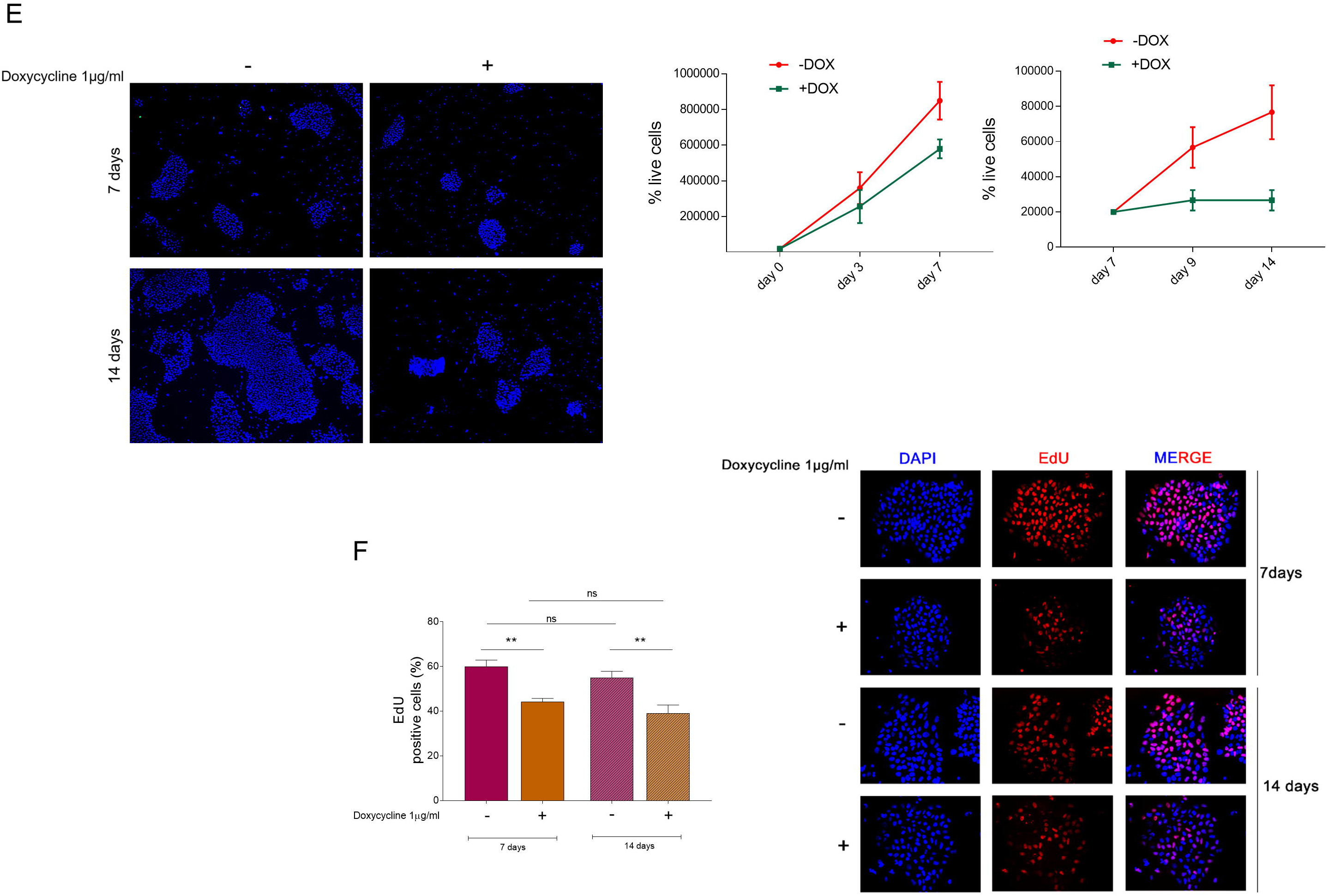
Generation and characterization of inducible SMARCAL1 knockdown iPSCs. A) Schematic representation of the experimental models. iPSCs derived from the human fibroblast were infected with the inducible RNAi lentivirus. When challenged with doxycycline 1µM for 7days, cells were considered SIOD. iPSCs were generated in absence of doxycycline and, once established, shifted in Dox+ media. B) Western blot showing the efficient silencing of SMARCAL1 after transduction of iPSCs with the Dox-inducible shSMARCAL1-containing lentiviral vector. Cells were tested by WB 72h after treatment with Dox. Lamin B1 was used as loading control. C) Western blotting shows the expression of SMARCAL1 upon shifting in Doxycycline-supplemented medium. WB was performed after 7 and 14 days of Doxycycline treatment. Lamin B1 was used as loading control. D) Comparative analysis by qRT-PCR of the expression of pluripotency markers NANOG and OCT4 after 7 days of doxycycline induction. Untreated iPSCs was used as reference sample. E) Analysis of iPSCs proliferation. Panel shows different size of DAPI-stained colonies from iPSCs cultured or not in the presence of doxycycline for the indicated time. Images are representative of different fields. The graphs show quantification of the total number of cells in each population after indicated periods of time in culture. Data are presented as means of three independent experiments. Horizontal black lines represent the mean ± SE. Student’s t test was used for statistical analyses. E) Analysis of replicating cells in SMARCAL1 depleted iPSCs. Replicating cells were labelled with EdU for 30min to mark S-phase and the graph plots the percentage of cells positive to EdU (red) at 7 and 14 days of doxycycline-induction. Representative images are shown, nuclear DNA was counterstained by DAPI (blue). Data are mean values from three independent experiments± standard error (SE). Statistical analysis was performed by ANOVA test, ns= not significant; **P ≤ 0.1.

Since the ultimate goal of the use of an iPSC model is differentiation in multiple cell types, we next analysed whether SMARCAL1 knockdown altered the expression of pluripotency genes. To this end, cells grown for 7 days in the presence or not of DOX were analysed for the expression levels of two key pluripotency genes (*NANOG and OCT4*) by real-time PCR. The analysis of gene expression showed that SMARCAL1 knockdown does not reduce the expression of the main pluripotency marker genes (Fig. 1D).

Having shown that continuous culturing in DOX-containing medium is effective in maintaining SMARCAL1 downregulated, we analysed whether depletion of SMARCAL1 affected proliferation in iSML iPSCs. To this end, iSML iPSCs were grown in the presence or not of DOX for 7 or 14 days and the number of live cells recorded over time. SMARCAL1 downregulation was able to greatly affect proliferation of iSML iPSCs (Fig. 1E). Strikingly, the effect on proliferation was particularly evident starting from 7 days of growth, as shown by the steady cell number and the reduced size of colonies (Fig. 1E). Consistently, iSML iPSCs cultured in the presence of DOX also showed a significant reduction in the number of replicating cells, as evidenced by the decreased number of EdU-positive cells, although no differences were observed between cells grown in DOX for 7 or 14 days (Fig. 1F). Notably, inducible depletion of SMARCAL1 induced reduced proliferation and EdU-incorporation also in normal human primary fibroblasts (Fig. S1A-C), suggesting that the phenotype is independent on the cell cycle type and not specific of iPSCs.

Collectively, these results indicate that inducible, long-term, depletion of SMARCAL1 in iPSCs is achievable. They also demonstrate that depletion of SMARCAL1, a condition mimicking the severe phenotype of SIOD cells, is sufficient to induce a time-dependent reduction in cell proliferation.

### Depletion of SMARCAL1 induces DNA damage and checkpoint activation in iSML iPSCs

Transformed or cancer-derived SMARCAL1-depleted cells are characterized by elevated levels of DNA damage (^5,12,13^). Since inducible SMARCAL1 downregulation hampers proliferation in iPSCs (Fig. 1), we analysed if this phenotype could correlate with enhanced DNA damage. To this end, we performed single-cell immunofluorescence analyses of the presence of two acknowledged markers of DNA damage and checkpoint activation, phosphorylated H2AX histone (γ-H2AX) and ATM (ATM-pSer1981). Depletion of SMARCAL1 by continuous cell growth in DOX resulted in a significant increase in the number of γ-H2AX-positive cells over time, which was otherwise not observed in cells cultured in the absence of DOX (Fig. 2A). Consistent with γ-H2AX data, depletion of SMARCAL1 also triggered ATM activation, as visualized by enhanced Ser1981 phosphorylation (Fig. 2B), an event associated with DNA damage and checkpoint activation. In contrast with γ-H2AX accumulation, the presence of ATM-pSer1981-positive cells was constant between 7 and 14 days of culture in DOX, while showed a small increase over time in iSML iPSCs growing in the absence of DOX (Fig. 2B). Increased activation of ATM and of downstream effectors such as CHK2 and KAP1 was investigated by Western blotting, confirming that depletion of iSML iPSCc cultured for 14 days in DOX show enhanced DNA damage and checkpoint activation (Fig. 2C). Of note, inducible depletion of SMARCAL1 in normal human primary fibroblasts also resulted in increased γ-H2AX and ATM-pSer1981 immunofluorescence at two different population doublings corresponding to a total of 14 days in culture with DOX, but the difference between the two population doublings tested was less striking than in iPSCs (Fig. S2A, B). Consistent with the role of SMARCAL1 as replication caretaker ^5,6^, culture of iSML1 iPSC in DOX-containing medium resulted in the large majority of S-phase cells staining positive for γ-H2AX and pATM foci, although staining was also observed in non-S-phase cells (Fig. 3A, B).

**Figure 2.**
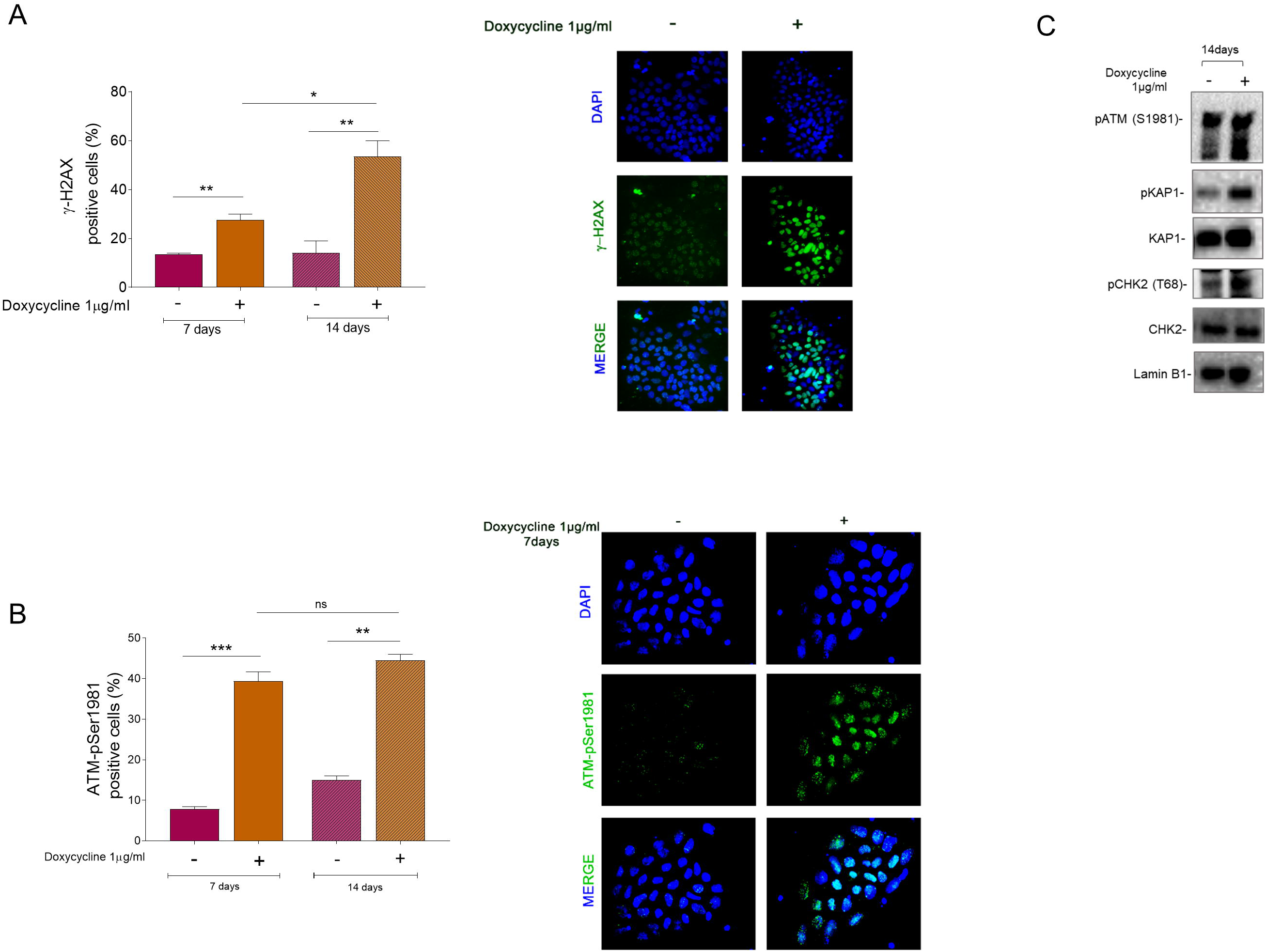
Depletion of SMARCAL1 induces DNA damage and checkpoint activation in iSML iPSCs. A) The iSML iPSCs were cultured for 7 and 14 days in the presence of doxycycline to induce SMARCAL1 downregulation and immunostained with anti-γ-H2AX antibody. The graph represents the analysis of the number of γ-H2AX positive cells Representative images are shown (γ-H2AX antibody (green), nuclear DNA, DAPI (blue)). Data are presented as mean± standard error (SE) from three independent experiments. *P<0.5; **P < 0.1, ANOVA test. B) The graph represents the analysis of the number of anti-ATM-pS1981 positive cells in iPSCs treated with Dox as indicated time Representative images are shown (ATM-pS1981 antibody (green), nuclear DNA, DAPI (blue)). Representative images from triplicate experiments are presented. Data are presented as mean ± standard error (SE) from three independent experiments. ns = not significant; **P < 0.1; ***P<0.01, ANOVA test. C) Immunoblot detection of the indicated DDR proteins in iSML iPSCs after 14 days of continuous treatment with doxycycline. Lamin B1 was used as the loading control protein.

**Figure 3.**
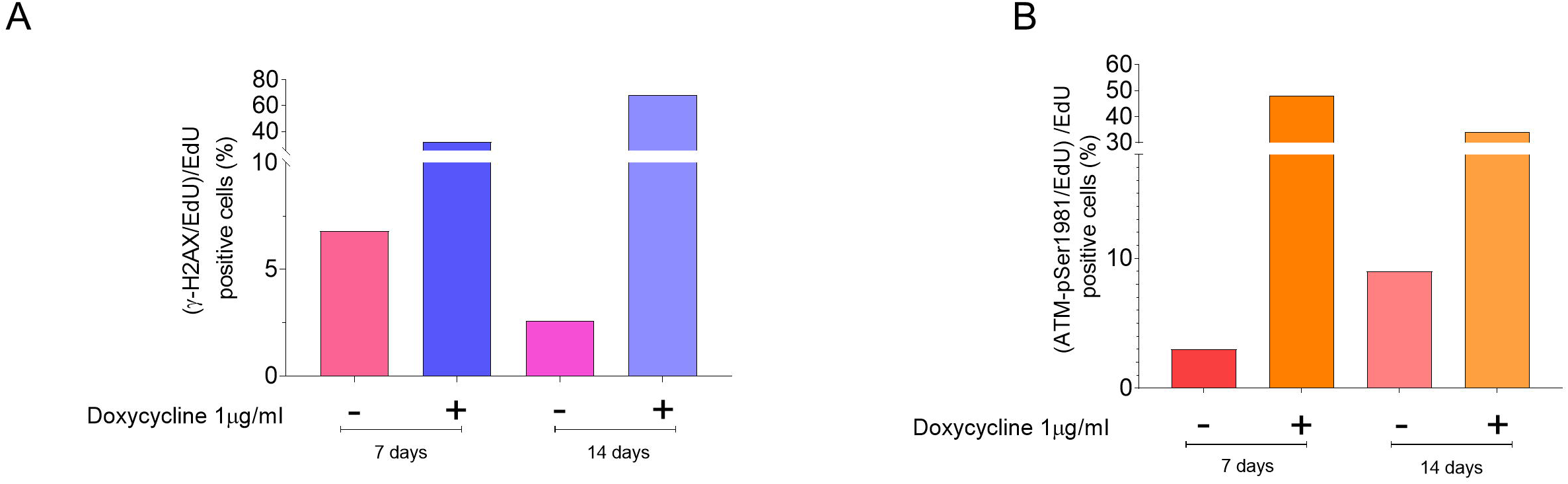
Enhanced phosphorylation of H2AX and ATM is related with S-phase. A) Analysis of the level of DNA damage in iSML iPSCs cells during replication. Cells were cultured in Dox+ medium as indicated and S-phase cells labelled for 30min before sampling. The graph reports the number of γ-H2AX positive cells in the S-phase population. B) Quantification of ATM-pS1981 staining in S-phase cells. The graph reports the number of γ-H2AX positive cells in the S-phase population. Data are from biological duplicates and are averages. Standard errors are not depicted and are < 15% of means.

These results indicate that continuous cell proliferation with reduced levels of SMARCAL1 leads to DNA damage accumulation. It also results in activation of proteins involved in the DNA-damage response, which is more striking in iPSCs than observed in primary fibroblasts.

### Depletion of SMARCAL1 in iPSCs resulted in reduced fork speed and defective replication

Having demonstrated that depletion of SMARCAL1 reduces proliferation of iPSCs cells and increases DNA damage, we tested if it also affected DNA replication dynamics. To this end, we performed single-molecule replication assays using dual-labelling with halogenated thymidine analogues and DNA fibres ^14^ (Fig. 4A). Analysis of IdU track length in dual-labelled fibres showed that loss of SMARCAL1 did not significantly reduce fork speed in iPSCs, although a small number of IdU tracks with reduced length was noticed in cells depleted of SMARCAL1 (Fig. 4B). Thus, we analysed the fork symmetry, another parameter linked to the presence of stalled forks ^15^, increasing labelling time to 30min (Fig. 4C). Notably, downregulation of SMARCAL1 resulted in an increasing number of asymmetric bi-directional forks, as evidenced by a Left/Right fork ratio higher than 1, where the left fork, per definition, is that showing the reduced length (Fig, 4C).

**Figure 4.**
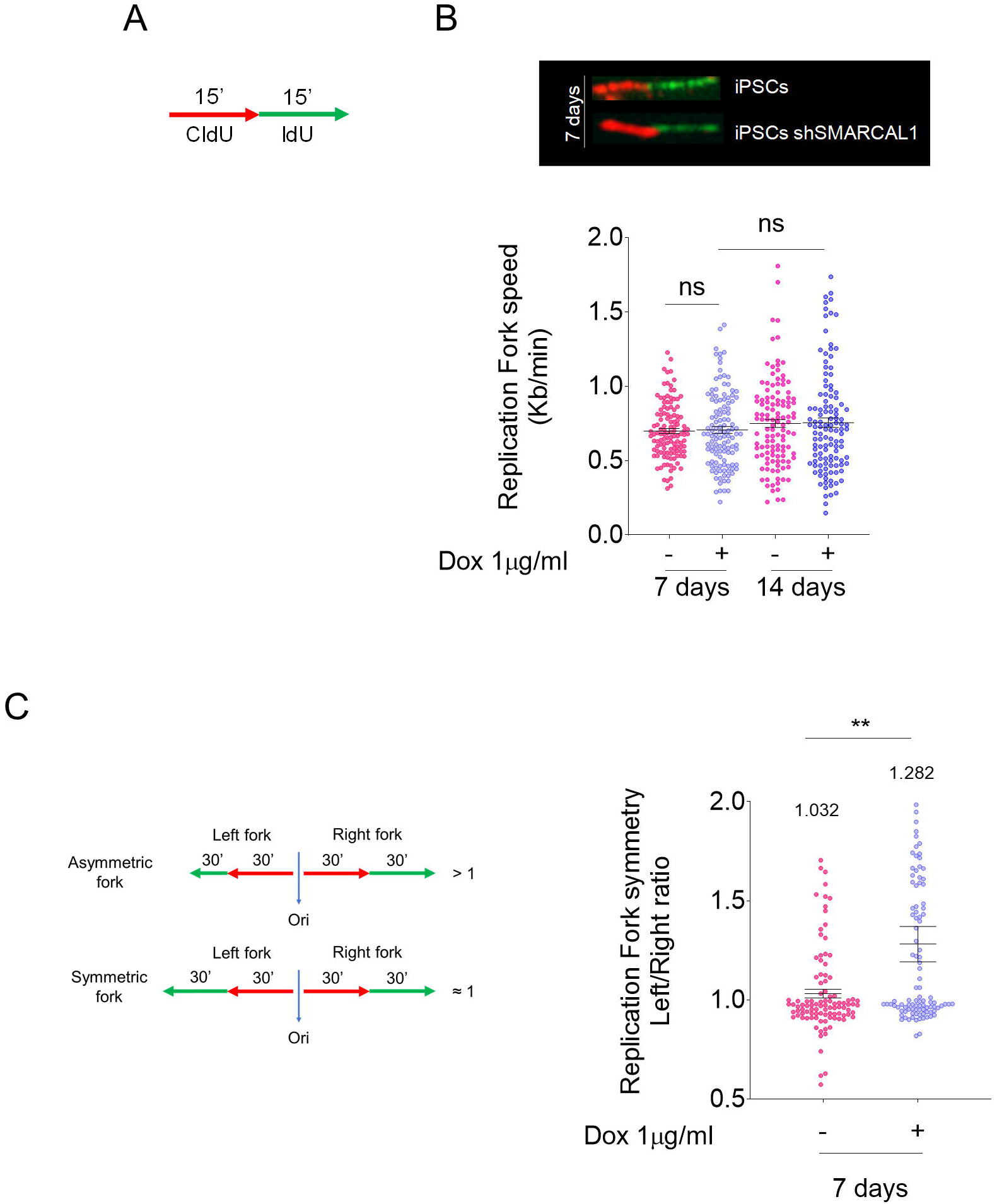
Analysis of DNA replication in SMARCAL1-depleted iPSCs. A) Schematic representation of the labelling strategy. Cells were labelled by two consecutive pulses of 15min with the indicated halogenated nucleotides. B) The length of the green, IdU, tracts were evaluated in at least 150 fibers and was converted in fork speed values. The graph shows scattered plot of single fork speed from iSML iPSCs treated as indicated. (NS, not significant; Mann-Whitney test). C) Analyses of the fork symmetry parameter evaluated as indicated in the cartoon. The graph shows the left/right fork value for each bi-directional fork (N=100 from two biological replicates). Values on top represents mean ratio for each population. (**, p<0.1; ANOVA test).

Collectively, these results indicate that downregulation of SMARCAL1 in iPSCs minimally affects DNA replication but induces a significant delay or stalling of a subset of replication forks.

### Preventing replication-transcription conflicts reduces DNA damage and replication defects in SMARCAL1-depleted iPSCs

Embryonic stem cells are characterised by reduced G1-phase ^16^, a condition reminiscent of cells with activated oncogenes and that is correlated with enhanced frequency of replication-transcription conflicts, which requires replication caretaker function ^17^. Hence, we tested whether increased DNA damage observed in iSML iPSCs grown in DOX was correlated with unresolved replication-transcription conflicts. To this end, we grew iSML iPSCs in DOX for 7 days and exposed cells to 5,6-dichloro-1-ß-d-ribofurosylbenzimidazole (DRB) in the last 4h before performing anti-γ-H2AX immunofluorescence. DRB is a transcription inhibitor and is widely used to prevent replication-transcription conflicts without affecting, in the short-term, proliferation ^18^, thus any reduction of γ-H2AX levels in cells treated with DRB is likely correlated with faulty resolution of accumulation of these events as demonstrated also un our group ^19^. Interestingly, while DRB treatment did not affect significantly the presence of γ-H2AX-positive cells in iSML iPSCs grown in the absence of DOX, it substantially reduced their number in cells depleted of SMARCAL1 resulting in a complete phenotype reversion (Fig. 5A). Consistent with the γ-H2AX data, DRB treatment determined also reversion of the enhanced ATM activation associated with SMARCAL1-depletion in iPSCs (Fig. 5B).

**Figure 5.**
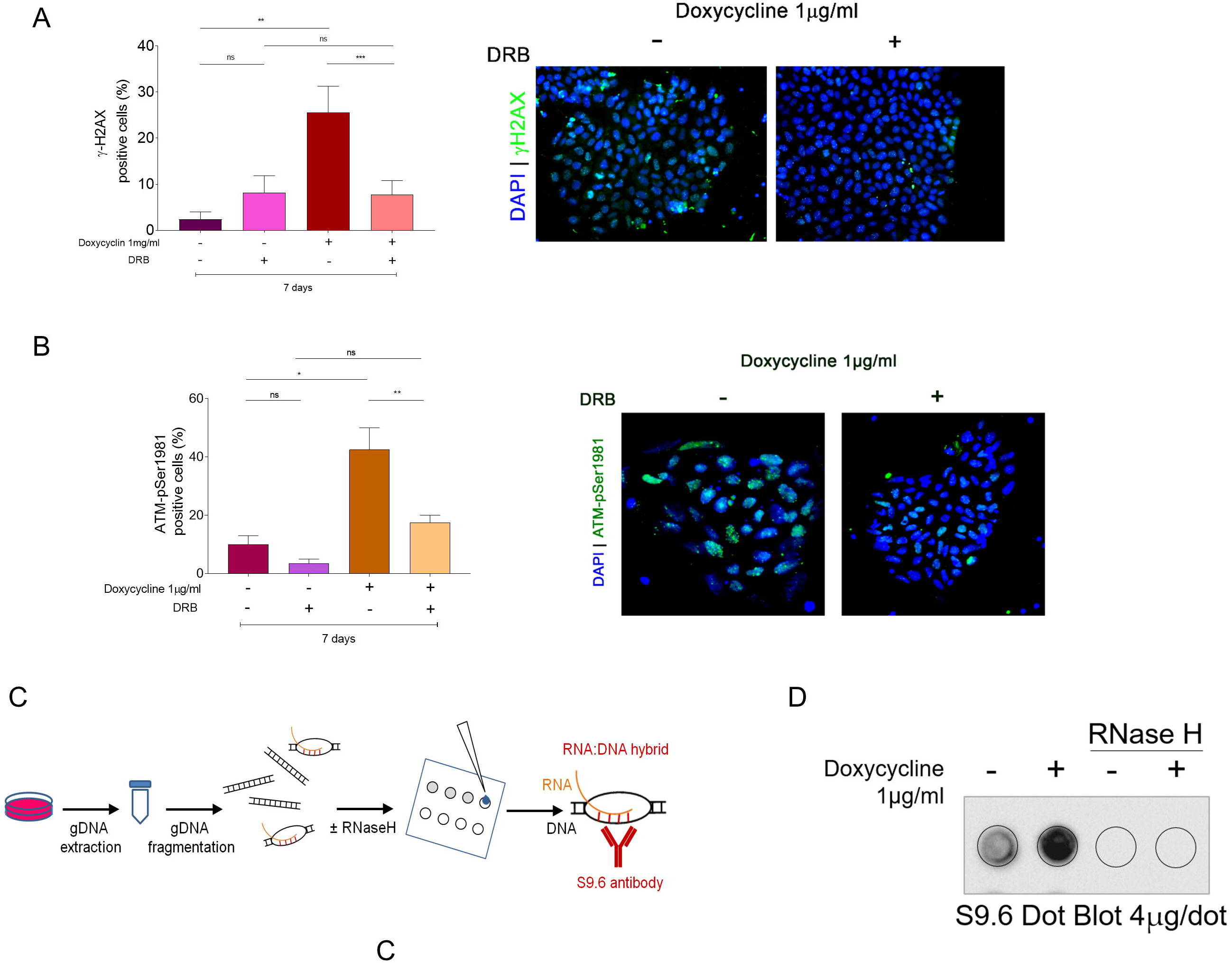
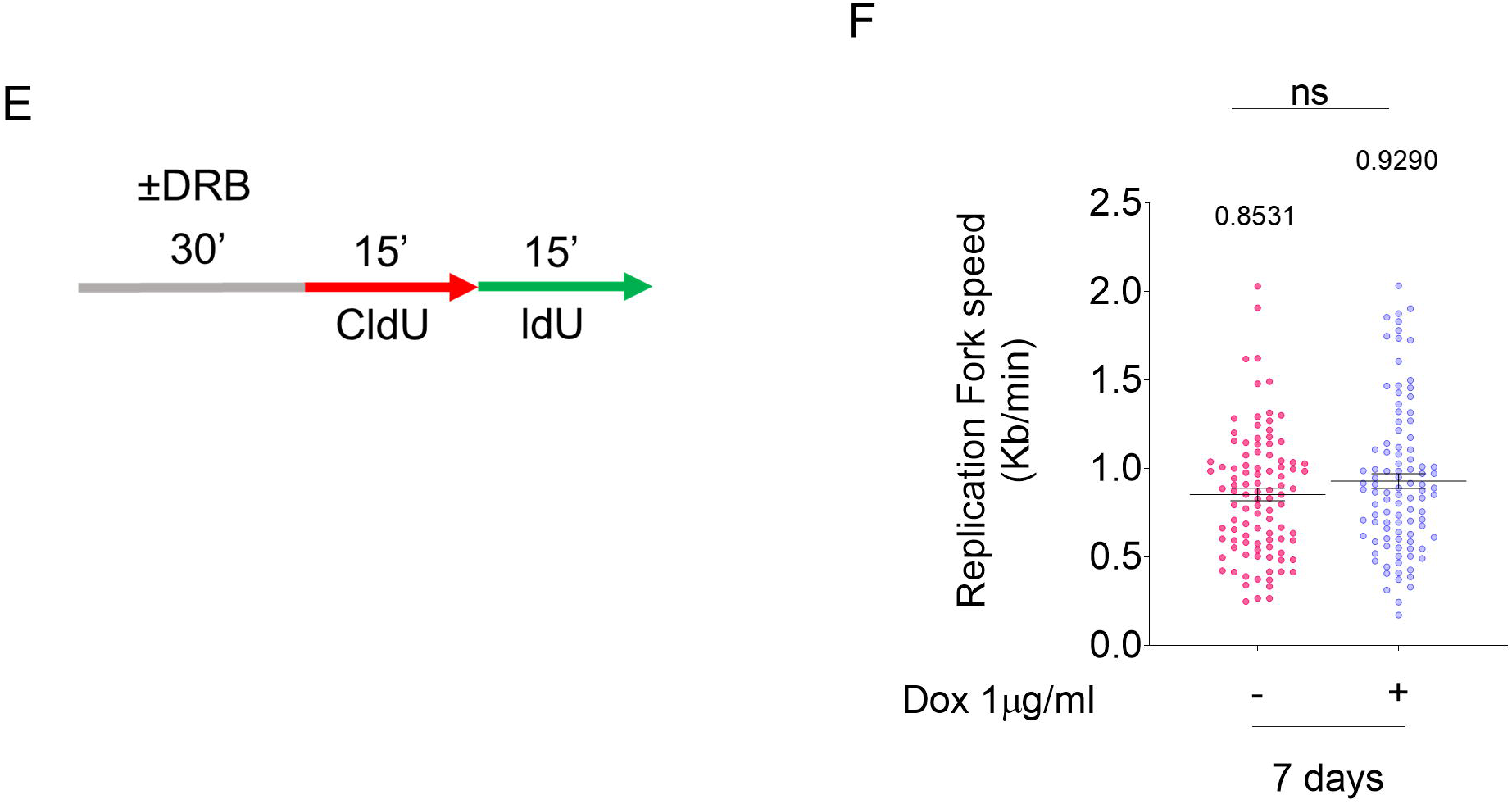
Preventing replication-transcription conflicts reduces DNA damage and DDR in SMARCAL1-depleted iPSCs. A) Analysis of accumulation of DNA damage in iSML iPSCs after SMARCAL1 downregulation. Four hours before sampling, DRB was added in the indicated samples at 50 µM. The graph shows the percentage of positive cells stained with anti γ-H2AX antibody. Representative images are shown. Nuclei were stained with DAPI (blue). Data are presented as mean± standard error (SE) from three independent experiments. NS= not significant; **P < 0.1; ***P <0.01, ANOVA test. B) SMARCAL1 depleted iPSCs were treated as in (A). Cells were stained with antiATM-pS1981 antibody (green) to detect phosphorylation of ATM and DDR activation. Representative images are presented in the right panel, nuclear DNA was counterstained by DAPI (blue). Data are presented as mean ± standard error (SE) from three independent experiments. NS = not significant; *P<0.5; **P < 0.1; ANOVA test. C) Experimental scheme for the detection of R-loops by dot blot in genomic DNA. D) Analysis of R-loop accumulation by dot blotting. Genomic DNA was isolated from iPSCs, treated or not with doxycycline to induce SMARCAL1 silencing, and then randomly fragmented prior to be spotted onto a nitrocellulose membrane. The control membrane was probed with anti-RNA–DNA hybrid S9.6 monoclonal antibody. Treatment with RNase H was used as a negative control. E) Schematic representation of the labelling strategy to detect replication fork progression. Cells were labelled by two consecutive pulses of 15min with the indicated halogenated nucleotides. DRB was added, where indicated, 4h before labelling. F) The length of the green, IdU, tracts were evaluated in at least 100 fibers and was converted in fork speed values. The graph shows scattered plot of single fork speed from iSML iPSCs treated as indicated. (NS, not significant; Mann-Whitney test).

Increased number of replication-transcription conflicts can be associated to enhanced accumulation of R-loops ^20,21^. To test if R-loops accumulated in iPSCs in which SMARCAL1 was downregulated, we purified genomic DNA from cells treated or not with DOX for 7 days and assessed the presence of R-loops by dot blot using the S9.6 anti-RNA-DNA hybrids antibody ^21,22^ (Fig. 5C). As shown in Figure 5D, cells depleted of SMARCAL1 had a substantially-elevated amount of genomic R-loop, suggesting that SMARCAL1 contributes to their prevention or resolution.

Accumulation of R-loops and replication-transcription conflicts may underlie DNA replication defects. Thus, we evaluated replication fork rate in iSML iPSCs cultured in DOX for 14 days and treated or not with DRB the last 4h (Fig. 5F). Interestingly, fork speed was unaffected by DRB treatment in SMARCAL1-depleted cells (+DOX).

Altogether, these results indicate that, in iPSCs growing in the absence of SMARCAL1, the increased DNA damage and depends on replication-transcription conflicts possibly deriving from accumulation of R-loops.

### Increased levels of replication-dependent DNA damage and checkpoint activation of SMARCAL1-depleted iPSCs persist upon their differentiation

Since we demonstrated that sustained depletion of SMARCAL1 induced the accumulation of replication defects and DNA damage in undifferentiated S-phase iSML iPSCs, we assessed if the presence of such DNA damage would persist also after spontaneous plurilineage differentiation.

To this end, iSML iPSCs were grown for 7 days in the presence of DOX before switching from pluripotency maintenance medium to differentiation conditions ^11^ (Fig. 6A). As shown in Figure 6B, SMARCAL1 knockdown was stable even in differentiated cells. Of note, the relative amount of SMARCAL1 in iSML iPSCs cultured without DOX (i.e. wild-type) declined during differentiation, consistent with its main function in replicating cells. We next analysed the presence of DNA damage by anti-γ-H2AX immunofluorescence in the population of differentiated iSML iPSCs. The number of cells staining positive for the DNA damage marker γ-H2AX was very limited in the population cultured in the absence of DOX, while it was much more elevated in cells grown under DOX (Figure 6C). Interestingly, immunofluorescence analysis of the activation of ATM, which is a readout of both DNA damage and checkpoint activation, revealed an even much more difference between cells cultured in the absence or presence of DOX (Fig. 6D). Notably, the percentage of actively-replicating cells was very low at the end of the 15days differentiation protocol, in both iSML iPSCs ±DOX, as evaluated by EdU incorporation (Fig. 6E). Thus, differences in the number of cells staining positive to the DNA damage markers are unlikely related to an excess of undifferentiated, replicating, cells in the iPSC growing in DOX.

**Figure 6.**
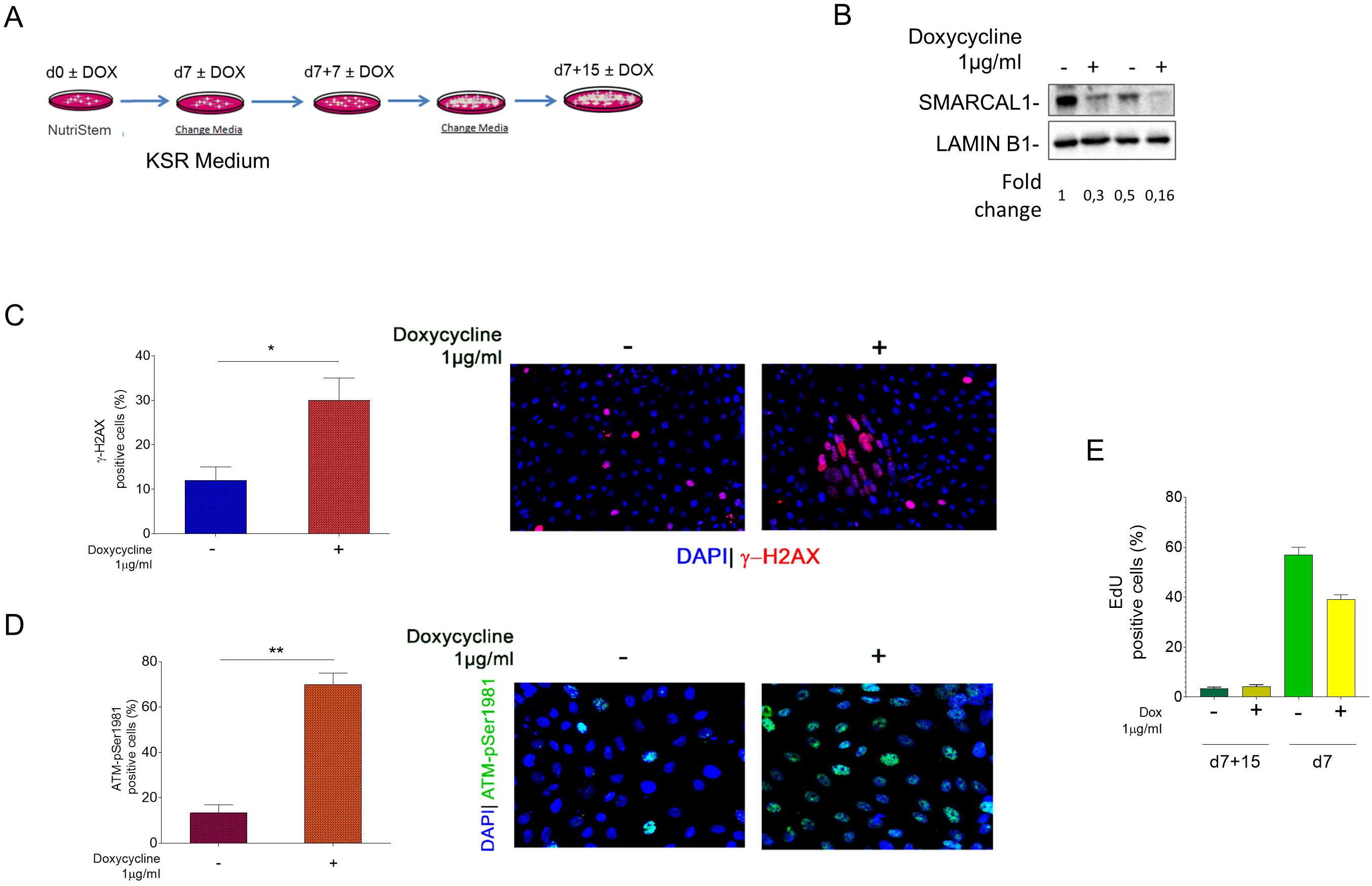
Increased levels of replication-dependent DNA damage and DDR activation persist in SMARCAL1-depleted iPSCs upon their differentiation. A) Scheme of the early differentiation protocol of iSML iPSCs. B) Western blot analysis of levels of SMARCAL1 depletion after 7 and 15 days from spontaneous multi-lineage differentiation. Lamin B1 was used as loading control. C-D) Analysis of DNA damage accumulation or DDR activation in iSML iPSCs after spontaneous multi-lineage differentiation. The graphs show the percentage of positive nuclei for each indicated endpoint. Data are presented as mean ± standard error (SE) from three independent experiments. *P<0.5, **p<0.1; ANOVA test. Representative images from cells stained with anti-γ-H2AX or anti-ATM-pS1981 antibody (red) are shown. Total nuclear DNA was counterstained by DAPI (blue). E) Analysis of replicating cells in SMARCAL1 depleted iPSCs. EdU labelling (30min) was used to mark replicating cells. The graph shows the percentage of cells positive to EdU signals (red) after 15 days of early differentiation treated or not with doxycycline. As reference, the values referred to the corresponding undifferentiated iSML iPSCs are included. Data are presented as mean ± standard error (SE) from three independent experiments.

Our data indicate that depletion of SMARCAL1 in iPSCs stimulates the accumulation of persistent DNA damage and long-term DNA damage response (DDR) activation also in differentiated cells. To determine whether such persistent DNA damage and active checkpoint signal would interfere with pluripotency, we induced formation of embryoid bodies and differentiation into the three germ layers and analysed expression of common marker genes by real-time PCR (Fig. 7A). Analyses of germ layer-specific genes showed that expression of Brachyury (mesoderm) Nestin (ectoderm) AFP and NR2F2 (an inhibitor of OCT4 expressed during early phases of human pluripotent stem cells differentiation; ^23^) were altered in cells depleted of SMARCAL1 while other genes were not affected (Fig. 7B).

**Figure 7.**
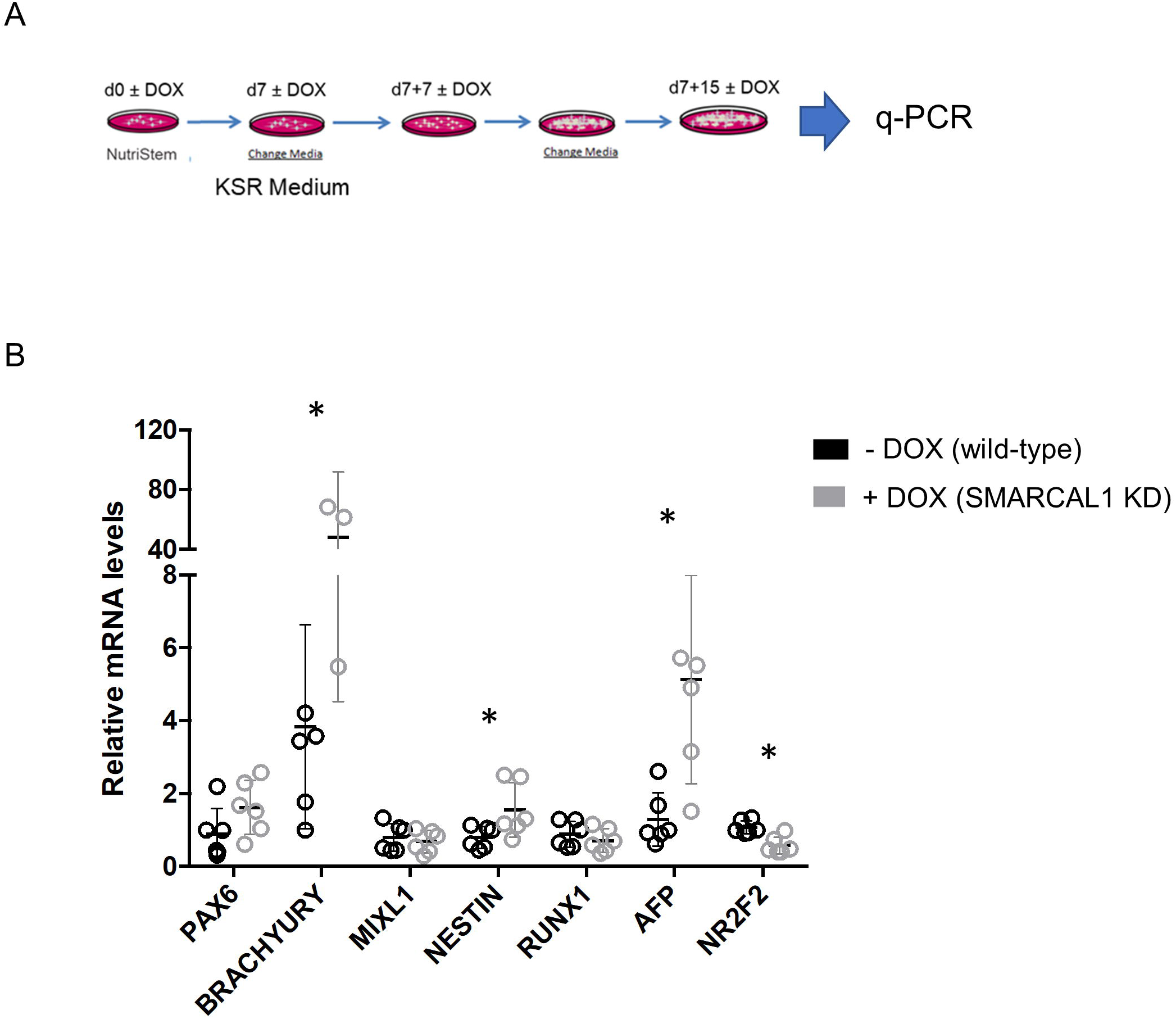
Depletion of SMARCAL1 affects expression of multi-lineage marker genes after differentiation of iPSCs. A) Scheme of the early differentiation protocol of iSML iPSCs. Gene expression was evaluated by q-PCR. B) Comparative analysis by q-PCR of the expression of the early differentiation markers shown in the graph in iPSCs treated as in (A). Untreated iPSCs was used as reference sample. Multiple t-test comparison (unpaired) was used to assess significance.

Collectively, these results indicate that accumulation of DNA damage and increased DDR caused by loss of SMARCAL1 in undifferentiated cells persist even after differentiation into the progenitors of the three germ-layers, suggesting that the effect of the loss of a replication caretaker maybe “inherited” by differentiating cells. Moreover, our data show that expression of germ layer-specific genes in iPSC-differentiated cells is affected by depletion of SMARCAL1.

## DISCUSSION

Here, we have shown that conditional knockdown of SMARCAL1 in human iPSCs induces replication-dependent and chronic accumulation of DNA damage stimulating activation of the DDR. Furthermore, our data indicate that accumulation of DNA damage and activation of the DNA damage response can be maintained in SMARCAL1-deficient iPSCs after differentiation possibly contributing to the altered expression of a subset of germ layer-specific master genes.

Mutations in *SMARCAL1* underlie SIOD ^4^, however how SMARCAL1 loss of function correlates with the disease phenotype is unknown. Depletion of SMARCAL1 in transformed cells has been shown to induce spontaneous DNA damage and proliferation defects, which are accrued by induced replication stress ^5,6,12^. Similarly, patient-derived transformed fibroblasts are characterized by DNA damage ^13^. Our inducible SMARCAL1 iPSCs appear to fully recapitulate these phenotypes showing the key cellular features of SMARCAL1 loss: spontaneous DNA damage and replication defects.

Of note, all animal models used to investigate on the SIOD pathogenesis failed to recapitulate the main disease phenotypes, with the exception of *Zebrafish* ^9,10^. Inducible PSCs are powerful models to study pathogenetic mechanisms of diseases ^24^. In this regard, our inducible SMARCAL1 knockdown iPSCs may prove useful tool for the identification of the molecular basis of SIOD and for genotype-phenotype correlation vis-à-vis of the reported involvement of additional environmental, genetic and/or epigenetic factors in the modulation of the penetrance of *SMARCAL1* mutations ^9,25^.

SMARCAL1 is a replication caretaker factor and its loss of function determines a mis-handling of perturbed replication forks ^12^. Furthermore, embryonic stem cells and iPSCs are characterized by spontaneous replication stress and by accumulation of remodelled stalled forks ^26^. SMARCAL1 is a crucial fork-remodelling protein ^27,28^. Hence, the function of SMARCAL1 is expected to be more important in iPSCs or in ESC then in other specialized cell types. Indeed, our data show that both DNA damage and activation of ATM increase over cell generation and mostly develop from S-phase cells. Consistent with this, proliferation potential of SMARCAL1 knockdown iPSCs does not decline immediately after depletion but shows a significant reduction after a week from the induced inactivation of the protein. Such a delayed kinetics suggests that DNA damage or replication stress should reach a threshold to induce proliferation arrest and is consistent with increased activation of ATM over time. Of note, our data indicate that conditional depletion of SMARCAL1 is sufficient to induce DNA damage, ATM activation and reduced proliferation in both primary fibroblasts and iPSCs. A telomeric function of SMARCAL1 has been also shown ^29^, however, the persistence of these phenotypes in iPSCs, which re-express hTERT ^30^, suggests that they are not specifically related with telomere erosion and supports the presence of a more genome-wide replication stress. Interestingly, persistence of phenotypes in iPSCs also differentiate SMARCAL1 loss from that of WRN, another critical replication caretaker ^31^. Indeed, proliferation potential of WS cells is rescued after reprogramming ^32^. From this point of view, SMARCAL1-depleted iPSCs behaves more likely as those generated from FA-A cells, which retain all the key cellular defects of the syndrome ^33^.

In pluripotent stem cells, the most likely source of replication stress is linked to short G1 phase and increased origin firing ^26,34^. A similar mechanism for the generation of replication stress has been put forward after oncogene activation ^17^. Of note, in this case, most of the replication stress would derive from interference between transcription and replication ^17^. Conditional knockdown of SMARCAL1 in iPSCs does trigger a substantial accumulation of R-loops, which are linked to transcription-replication conflicts ^20,21^. This observation would suggest that SMARCAL1 counteracts accumulation of R-loops and transcription-replication conflicts as supported by rescue of DNA damage and ATM activation by transcription inhibition (Fig 5).

The recent observation of a non-canonical ATM activation that is dependent on R-loop accumulation and alternative processing ^35^, and increased upon defective replication or mild replication stress ^19^, is consistent with our data. Notably, although *Smarcal1* KO MEFs do not have any significant proliferation defect compared with wild-type cells, they show a slow-growth phenotype if treated with α-amanitin ^9^. Since α-amanitin interferes with elongation phase of transcription and not with initiation like DRB, it is possible that slowed RNA polII increases the chance of replication-transcription conflicts in *Smarcal1* KO MEFs, resulting in a proliferation defect as observed in our iPSC model.

Interestingly, DNA damage and ATM activation caused by transcription-replication interference in iPSCs depleted of SMARCAL1 persist also after differentiation of embryonic body in cells of the three germ layers. Thus, a replication-dependent phenotype seems to be inherited in differentiated cells. Most importantly, loss of SMARCAL1 affects expression of a subset of germ layer-specific master genes. The pathogenetic mechanisms responsible for SIOD are still elusive, however, SMARCAL1 deficiency has been reported to pathologically modulate gene expression ^9,25,36–38^. An intriguing possibility is that loss of SMARCAL1 function indirectly influences gene expression through increased levels of replication stress, as suggested for loss of WRN or FANCJ helicase ^39–41^. Accumulation or persistence of R-loops could be involved in this mechanism making attractive to assess if R-loops preferentially accumulates at affected genes.

One of the germ layer master genes showing increased expression in SMARCAL1 knockdown iPSCs is *BRACHYURY*. Notably, expression of *BRACHYURY* has been found elevated in cordomas and correlates with increased cellular proliferation in the bone ^42^, hence providing a possible link to osseous dysplasia, which is one of the clinical phenotypes of SIOD ^3^.

Altogether, our work indicates that conditional downregulation of SMARCAL1 in iPSCs recapitulates phenotypes observed in specialized cells following SMARCAL1 depletion. Most importantly, our study demonstrates that loss of SMARCAL1 induces the accumulation of DNA damage and ATM activation in iPSCs through replication stress correlated with replication-transcription conflicts. Since mutations in SMARCAL1 cause the multisystemic genetic disease SIOD^4^ and complete loss of function of SMARCAL1 correlates with the severest form of this conditions ^7^, our conditional knockdown of SMARCAL1 in iPSCs may represent a powerful model where studying pathogenetic mechanisms overcoming the reported inability of other model systems to recapitulate the disease.

## ACKNOWLEDGMENTS

This work was supported by Fondazione Telethon (GEP15050 to PP) and Fondazione Terzo Pilastro Internazionale (to PP), and in part by Associazione Italiana per la Ricerca sul Cancro (AIRC IG15410 to AF) and by ARISLA (to AR).

## AUTHOR CONTRIBUTIONS

G.M.P. performed all cellular analysis in iPSCs. F.S. contributed to iPSCs maintenance, analysis of proliferation and to gene expression analysis. V.P. performed lentiviral transduction and the analysis of replication fork progression. V.M. performed the analysis of R-loop. N.M. contributed to the generation of inducible primary fibroblasts and performed analysis in primary fibroblasts. A.R. contributed to iPSCs generation and supervised maintenance of iPSCs. P.P. contributed to replication fork symmetry analysis. G.M.P., F.S. V.M and V.P. analysed data and contributed to designing the experiments and writing the paper. A.R., A.F. and P.P designed experiments, analysed data and wrote the paper. All authors approved the paper.

## MATERIALS AND METHODS

### Human iPSC culture, infection and differentiation

Human iPSCs used in this study belong to the WT I line described in Lenzi et al., 2015 ^11^. iSML iPSCs were generated by spinfection of iPSCs (passage 13) with the lentiviral vector Tet-ON-shSMARCAL1 at 0.5 of MOI. Three days later, infected cells were selected with 1 μg/mL puromycin and maintained under selection for 4 days. When indicated, 1 μM doxycycline was added to the medium to induce shSMARCAL1 expression. Established iSML iPSCs were maintained in Nutristem-XF (Biological Industries) in plates coated with hESC-qualified matrigel (BD Biosciences) and passaged every 4-5 days with 1 mg/ml dispase (Gibco).

For spontaneous plurilineage differentiation, 48 h after passaging the culture medium was changed to KSR Medium (DMEM-F12, Sigma-Aldrich; 15% Knockout Serum Replacement, Thermo Fisher Scientific; 1X Glutamax, Thermo Fisher Scientific; 1X Non-Essential Aminoacids, Thermo Fisher Scientific; 100 U/ml Penicillin + 100 μg/ml Streptomycin, Sigma-Aldrich). Medium was refreshed every other day until the end of differentiation.

### Real-time qRT-PCR analysis of differentiation markers

Total RNA was extracted with the Quick RNA MiniPrep (Zymo Research) and retrotranscribed with PrimeScript RT reagent Kit (Perfect Real Time). Targets were analyzed by real-time qRT-PCR with SYBR Green Power-UP (Thermo Fisher Scientific) in a 7500 Fast Real Time PCR System (Thermo Fisher Scientific) and calculations performed with the delta delta Ct method. The internal control used was the housekeeping gene ATP5O. Primer sequences are reported in Lenzi et al., ^11^.

### Growth curve

The cells were seeded at 1.8 × 104 cells per plate. After trypsinization, cells were counted through electronic counting cells (BioRad) for the following 6 days. The growth curve of the cell cultures was expressed as number of cells as a function of time.

### DNA fibre analysis

Cells were pulse-labelled with 25 μM CldU and then labeled with 250 μM IdU with or without treatment as reported in the experimental schemes. DNA fibers were prepared and spread out as previously described ^43^. Images were acquired randomly from fields with untangled fibres using Eclipse 80i Nikon Fluorescence Microscope, equipped with a Video Confocal (ViCo) system. A minimum of 100 individual fibres were analyzed for each experiment and each experiment was repeated three times.

### Western blot analysis

Western blots were performed using standard methods. The antibodies used are listed below. Blots were developed using Westernbright ECL (Advasta) according to the manufacturer’s instructions. Quantification was performed on scanned images of blots using Image Lab software, and values shown on the graphs represent a normalization of the protein content evaluated through LaminB1.

### Antibodies

The primary antibodies used were: anti-SMARCAL1 (1:1000; abcam), anti-pCHK2 (1:1000; Cell Signaling), anti-CHK2 (1:1000; Cell Signaling), anti-pKAP1 (1.1000; Cell Signaling), anti-KAP1 (1:1000; Cell Signaling), anti p-ATM (1:800; Cell Signaling), anti-pATM (IF 1:300; Millipore), anti-ATM (1.1000; Novus), anti-pS139H2A.X (1:1000; Millipore), anti-LaminB1 (1:20000; abcam). HRP-conjugated matched secondary antibodies were from Jackson Immunoresearch and were used at 1:40000.

### Immunofluorescence

Immunofluorescence microscopy was performed on cells grown on cover-slips. Briefly, cells were fix with 4% PFA and permeabilized with 0,4%Triton-X 100/PBS. After blocking, coverslips were incubated for 1h at RT with the indicated antibodies. For detection of anti-BrdU, after permeabilization with 0,4%Triton-X 100/PBS, cells were denatured in HCl 2.5N for 45 min at RT. Alexa Fluor® 488 conjugated-goat anti mouse and Alexa Fluor® 594 conjugated-goat anti-rabbit secondary antibodies (Life Technologies) were used at 1:200. Nuclei were stained with 4’,6-diamidino-2-phenylindole (DAPI 1:4000, Serva). Coverslips were observed at 40× objective with the Eclipse 80i Nikon Fluorescence Microscope, equipped with a Video-Confocal (ViCo) system. Images were processed by using Photoshop (Adobe) program to adjust contrast and brightness. For each time point at least 200 nuclei were examined. Parallel samples incubated with either the appropriate normal serum or only with the secondary antibody confirmed that the observed fluorescence pattern was not attributable to artefacts. Experiments for labelling cellular DNA with EdU (5-ethynyl-2’-deoxyuridine). EdU was added to the culture media (10 µM), for 30 min. Detection of EdU was performed used Click-iT EdU imaging Kits (Invitrogen for green signal and Base Click for red signal).

### Dot Blot Analysis

Dot blot analysis was performed according to the protocol reported elsewhere ^44^. Genomic DNA was isolated by standard extraction with phenol/clorophorm/isoamylic alcohol (pH 8.0) followed by precipitation with 3 M NaOAc and 70% ethanol. Isolated gDNA was randomly fragmented overnight at 37°C with a cocktail of restriction enzymes (BsrgI, EcoRI, HindIII, XbaI) supplemented with 1 M Spermidin. After incubation, digested DNA was cleaned up with phenol/chloroform extraction and standard Ethanol precipitation. After sample quantification, 5 μg of digested DNA were incubated with RNase H overnight at 37°C as a negative control. Five micrograms of each sample were spotted onto a nitrocellulose membrane, blocked in 5% non-fat dry milk and incubated with the anti-DNA–RNA hybrid [S9.6] antibody (Kerafast) overnight at 4°C. Horseradish peroxidase-conjugated goat specie-specific secondary antibody (Santa Cruz Biotechnology, Inc.) was used. Quantification on scanned image of blot was performed using Image Lab software.

## REFERENCES

1. Boerkoel CF, O’Neill S, André JL, Benke PJ, Bogdanovíc R, Bulla M et al. Manifestations and treatment of Schimke immuno-osseous dysplasia: 14 new cases and a review of the literature. Eur J Pediatr 2000; 159: 1–7.

2. Saraiva JM, Dinis A, Resende C, Faria E, Gomes C, Correia AJ et al. Schimke immuno-osseous dysplasia: case report and review of 25 patients. J Med Genet 1999; 36: 786–9.

3. Clewing JM, Antalfy BC, Lücke T, Najafian B, Marwedel KM, Hori A et al. Schimke immuno-osseous dysplasia: a clinicopathological correlation. J Med Genet 2007; 44: 122–30.

4. Boerkoel CF, Takashima H, John J, Yan J, Stankiewicz P, Rosenbarker L et al. Mutant chromatin remodeling protein SMARCAL1 causes Schimke immuno-osseous dysplasia. Nat Genet 2002; 30: 215–20.

5. Ciccia A, Bredemeyer AL, Sowa ME, Terret M-E, Jallepalli P V., Harper JW et al. The SIOD disorder protein SMARCAL1 is an RPA-interacting protein involved in replication fork restart. Genes Dev 2009; 23: 2415–2425.

6. Bansbach CE, Bétous R, Lovejoy C a., Glick GG, Cortez D. The annealing helicase SMARCAL1 maintains genome integrity at stalled replication forks. Genes Dev 2009; 23: 2405–2414.

7. Elizondo LI, Cho KS, Zhang W, Yan J, Huang C, Huang Y et al. Schimke immuno-osseous dysplasia: SMARCAL1 loss-of-function and phenotypic correlation. J Med Genet 2009; 46: 49–59.

8. Elizondo LI, Huang C, Northrop JL, Deguchi K, Clewing JM, Armstrong DL et al. Schimke immuno-osseous dysplasia: a cell autonomous disorder? Am J Med Genet A 2006; 140: 340– 8.

9. Baradaran-Heravi A, Cho KS, Tolhuis B, Sanyal M, Morozova O, Morimoto M et al. Penetrance of biallelic SMARCAL1 mutations is associated with environmental and genetic disturbances of gene expression. Hum Mol Genet 2012; 21: 2572–87.

10. Huang C, Gu S, Yu P, Yu F, Feng C, Gao N et al. Deficiency of smarcal1 causes cell cycle arrest and developmental abnormalities in zebrafish. Dev Biol 2010; 339: 89–100.

11. Lenzi J, De Santis R, de Turris V, Morlando M, Laneve P, Calvo A et al. ALS mutant FUS proteins are recruited into stress granules in induced pluripotent stem cell-derived motoneurons. Dis Model Mech 2015; 8: 755–66.

12. Couch FB, Bansbach CE, Driscoll R, Luzwick JW, Glick GG, Bétous R et al. ATR phosphorylates SMARCAL1 to prevent replication fork collapse. Genes Dev 2013; 27: 1610–1623.

13. Bansbach CE, Boerkoel CF, Cortez D. SMARCAL1 and replication stress: An explanation for SIOD? Nucleus 2010; 1: 245–248.

14. Leuzzi G, Marabitti V, Pichierri P, Franchitto A. WRNIP1 protects stalled forks from degradation and promotes fork restart after replication stress. EMBO J 2016; 1–15.

15. Técher H, Koundrioukoff S, Azar D, Wilhelm T, Carignon S, Brison O et al. Replication dynamics: Biases and robustness of DNA fiber analysis. J Mol Biol 2013; 425: 4845–4855.

16. White J, Dalton S. Cell cycle control of embryonic stem cells. Stem Cell Rev 2005; 1: 131–8.

17. Macheret M, Halazonetis TD. Intragenic origins due to short G1 phases underlie oncogeneinduced DNA replication stress. Nature 2018; 555: 112–116.

18. Salas-Armenteros I, Pérez-Calero C, Bayona-Feliu A, Tumini E, Luna R, Aguilera A. Human THO–Sin3A interaction reveals new mechanisms to prevent R-loops that cause genome instability. EMBO J 2017; 36: 3532–3547.

19. Marabitti V, Lillo G, Malacaria E, Palermo V, Sanchez M, Pichierri P et al. ATM pathway activation limits R-loop-associated genomic instability in Werner syndrome cells. Nucleic Acids Res 2019. doi:10.1093/nar/gkz025.

20. Lang KS, Hall AN, Merrikh CN, Ragheb M, Tabakh H, Pollock AJ et al. Replication-Transcription Conflicts Generate R-Loops that Orchestrate Bacterial Stress Survival and Pathogenesis. Cell 2017; 170: 787–799.e18.

21. Hamperl S, Bocek MJ, Saldivar JC, Swigut T, Cimprich KA. Transcription-Replication Conflict Orientation Modulates R-Loop Levels and Activates Distinct DNA Damage Responses. Cell 2017; 170: 774–786.e19.

22. Boguslawski SJ, Smith DE, Michalak MA, Mickelson KE, Yehle CO, Patterson WL et al. Characterization of monoclonal antibody to DNA.RNA and its application to immunodetection of hybrids. J Immunol Methods 1986; 89: 123–30.

23. Rosa A, Brivanlou AH. A regulatory circuitry comprised of miR-302 and the transcription factors OCT4 and NR2F2 regulates human embryonic stem cell differentiation. EMBO J 2011; 30: 237–248.

24. Avior Y, Sagi I, Benvenisty N. Pluripotent stem cells in disease modelling and drug discovery. Nat Rev Mol Cell Biol 2016; 17: 170–182.

25. Morimoto M, Choi K, Boerkoel CF, Cho KS. Chromatin changes in SMARCAL1 deficiency: A hypothesis for the gene expression alterations of Schimke immuno-osseous dysplasia. Nucleus 2016; 7: 560–571.

26. Ahuja AK, Jodkowska K, Teloni F, Bizard AH, Zellweger R, Herrador R et al. A short G1 phase imposes constitutive replication stress and extensive fork remodeling in mouse embryonic stem cells. Nat Commun 2016; In press: 1–11.

27. Bétous R, Mason AC, Rambo RP, Bansbach CE, Badu-Nkansah A, Sirbu BM et al. SMARCAL1 catalyzes fork regression and holliday junction migration to maintain genome stability during DNA replication. Genes Dev 2012; 26: 151–162.

28. Kolinjivadi AM, Sannino V, De Antoni A, Zadorozhny K, Kilkenny M, Técher H et al. Smarcal1-Mediated Fork Reversal Triggers Mre11-Dependent Degradation of Nascent DNA in the Absence of Brca2 and Stable Rad51 Nucleofilaments. Mol Cell 2017; 67: 867–881.e7.

29. Poole LA, Zhao R, Glick GG, Lovejoy CA, Eischen CM, Cortez D. SMARCAL1 maintains telomere integrity during DNA replication. Proc Natl Acad Sci U S A 2015; 112: 14864– 14869.

30. Takahashi K, Tanabe K, Ohnuki M, Narita M, Ichisaka T, Tomoda K et al. Induction of Pluripotent Stem Cells from Adult Human Fibroblasts by Defined Factors. Cell 2007; 131: 861–872.

31. Franchitto A, Pichierri P. Replication fork recovery and regulation of common fragile sites stability. Cell Mol Life Sci 2014; 71: 4507–4517.

32. Shimamoto A, Kagawa H, Zensho K, Sera Y, Kazuki Y, Osaki M et al. Reprogramming Suppresses Premature Senescence Phenotypes of Werner Syndrome Cells and Maintains Chromosomal Stability over Long-Term Culture. PLoS One 2014; 9: e112900.

33. Liu G-H, Suzuki K, Li M, Qu J, Montserrat N, Tarantino C et al. Modelling Fanconi anemia pathogenesis and therapeutics using integration-free patient-derived iPSCs. Nat Commun 2014; 5: 4330.

34. Ryba T, Hiratani I, Lu J, Itoh M, Kulik M, Zhang J et al. Evolutionarily conserved replication timing profiles predict long-range chromatin interactions and distinguish closely related cell types. Genome Res 2010; 20: 761–770.

35. Tresini M, Warmerdam DO, Kolovos P, Snijder L, Vrouwe MG, Demmers JAA et al. The core spliceosome as target and effector of non-canonical ATM signalling. Nature 2015; 523: 53–8.

36. Morimoto M, Myung C, Beirnes K, Choi K, Asakura Y, Bokenkamp A et al. Increased Wnt and Notch signaling: a clue to the renal disease in Schimke immuno-osseous dysplasia? Orphanet J Rare Dis 2016; 11: 149.

37. Sanyal M, Morimoto M, Baradaran-Heravi A, Choi K, Kambham N, Jensen K et al. Lack of IL7Rα expression in T cells is a hallmark of T-cell immunodeficiency in Schimke immuno-osseous dysplasia (SIOD). Clin Immunol 2015; 161: 355–365.

38. Morimoto M, Wang KJ, Yu Z, Gormley AK, Parham D, Bogdanovic R et al. Transcriptional and posttranscriptional mechanisms contribute to the dysregulation of elastogenesis in Schimke immuno-osseous dysplasia. Pediatr Res 2015; 78: 609–617.

39. Schiavone D, Guilbaud G, Murat P, Papadopoulou C, Sarkies P, Prioleau M-N et al. Determinants of G quadruplex-induced epigenetic instability in REV1-deficient cells. EMBO J 2014; 33: 2507–20.

40. Papadopoulou C, Guilbaud G, Schiavone D, Sale JE. Nucleotide Pool Depletion Induces G- Quadruplex-Dependent Perturbation of Gene Expression. Cell Rep 2015; 13: 2491–2503.

41. Khurana S, Oberdoerffer P. Replication Stress: A Lifetime of Epigenetic Change. Genes (Basel) 2015; 6: 858–77.

42. Miettinen M, Wang Z, Lasota J, Heery C, Schlom J, Palena C. Nuclear Brachyury Expression Is Consistent in Chordoma, Common in Germ Cell Tumors and Small Cell Carcinomas, and Rare in Other Carcinomas and Sarcomas. Am J Surg Pathol 2015; 39: 1305–1312.

43. Iannascoli C, Palermo V, Murfuni I, Franchitto A, Pichierri P. The WRN exonuclease domain protects nascent strands from pathological MRE11/EXO1-dependent degradation. Nucleic Acids Res 2015; 43: 9788–803.

44. Morales JC, Richard P, Patidar PL, Motea EA, Dang TT, Manley JL et al. XRN2 Links Transcription Termination to DNA Damage and Replication Stress. PLoS Genet 2016; 12: e1006107.

