## Supplementary figures 1 and 2, and legends for "Generation of inducible SMARCAL1 knock-down iPSC to model severe Schimke immune-osseous dysplasia reveals a link between replication stress and altered expression of master differentiation genes"

Giusj Monia Pugliese et al.

SUPPLEMENTARY FIGURE LEGENDS

**Figure S1. Depletion of SMARCAL1 induced reduced proliferation in normal human primary fibroblasts**

A) Western blot showing SMARCAL1 depletion in primary fibroblasts after switching in Dox+ medium. Doxycycline was added a p14 and analysis was performed at p17 and p22. Lamin B1 is used for normalization. B) Evaluation of cell population size for wild-type and shSMARCAL1-induced primary fibroblasts. A starting culture of 6x10^4^ cells was used to plate identical numbers of cells for each cell line and after 5 days in culture the total number of cells was recorded and reported in graph. Data are means±SE from two independent experiments. (**, p<0.1; ANOVA test). C) Evaluation of the proliferating population in wild-type and shSMARCAL1-induced primary fibroblasts. Doxycycline was added a p14 and analysis was performed at p17 and p22. Cells were cultured in IdU-containing medium in the last 24h before analysis. Graph shows the number of IdU+. Data are from biological duplicates and are averages. Standard errors are not depicted and are < 15% of means. Representative images of the immunofluorescence experiment are shown. Total DNA is stained with DAPI.

**Figure S2. SMARCAL1-silenced primary fibroblast shows increased DNA damage and DDR activation**

A-B) Analysis of spontaneous DNA damage and DDR in primary fibroblast depleted for SMARCAL1. Doxycycline was added a p14 and analysis was performed at p17 and p22. Cells were immunostained with anti-γ-H2AX or anti-ATM-pS1981 antibody. The graphs represent the analysis of positive cells after continuous treatment with doxycycline at p17 and p22 (i.e. 7 and 12 days). Representative images of fluorescence fields from cells stained with anti-γ-H2AX or anti-ATM-pS1981 antibody (green) are provided. Total nuclear DNA was counterstained by DAPI (blue). Data are from biological duplicates and are averages. Standard errors are not depicted and are < 15% of means.

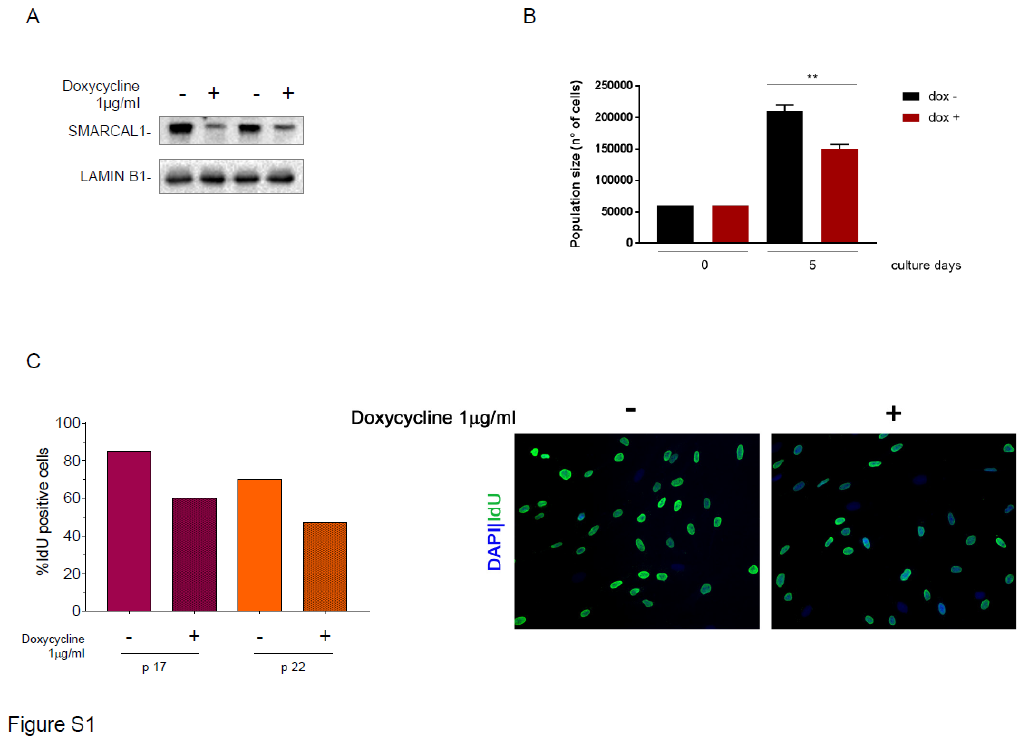
SUPPLEMENTARY FIGURES

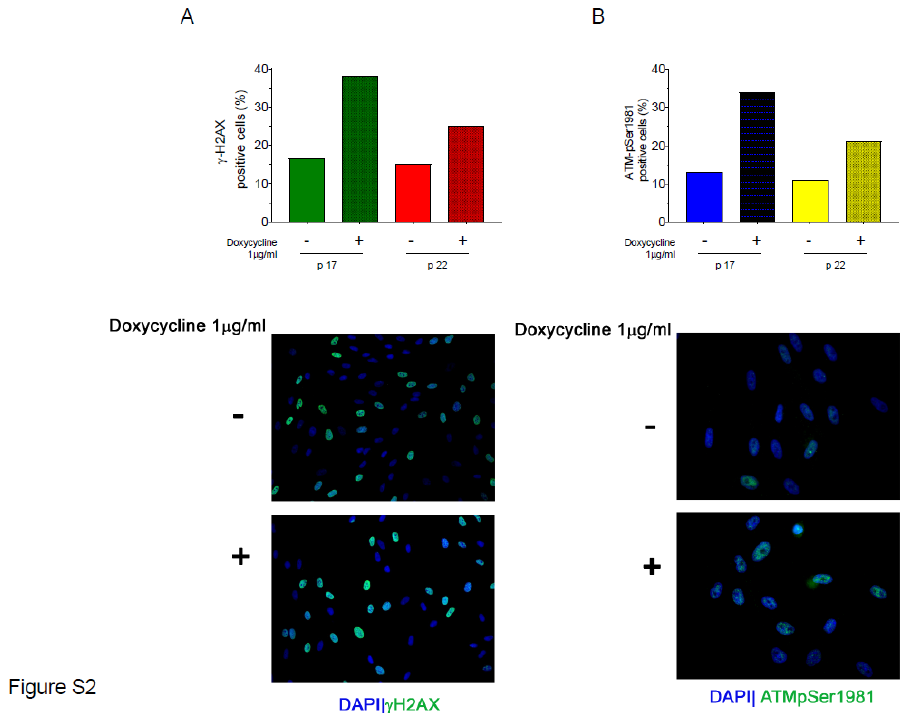
